# Prey-Driven Behavioral Habitat Use in a Low-Energy Ambush Predator

**DOI:** 10.1101/2020.09.17.301697

**Authors:** Annalee M. Tutterow, Andrew S. Hoffman, John L. Buffington, Zachary T. Truelock, William E. Peterman

**Affiliations:** School of Environment and Natural Resources, The Ohio State University, 2021 Coffey Rd, 210 Kottman Hall, Columbus, OH 43210, USA

**Keywords:** Ambush hunting, Camera traps, Habitat use, Optimal foraging, Prey availability, Site selection, Small mammals, Timber rattlesnakes

## Abstract

1. Food acquisition is an important modulator of animal behavior and habitat selection that can affect fitness. Optimal foraging theory predicts that predators should select habitat patches to maximize their foraging success and net energy gain, which predators can achieve by targeting spaces with high prey availability. However, it is debated whether prey availability drives fine-scale habitat selection for predators.
2. We assessed whether an ambush predator, the timber rattlesnake (*Crotalus horridus*), exhibits optimal foraging site selection based on the spatial distribution and availability of prey.
3. We evaluated the spatial concordance of radio-telemetered timber rattlesnake foraging locations and passive infrared game camera trap detections of potential small mammal prey (*Peromyscus* spp., *Tamias striatus,* and *Sciurus* spp.) in a mixed-use forest in southeastern Ohio from 2016–2019. We replicated a characteristic timber rattlesnake ambush position by focusing cameras over logs and modeled small mammal encounters across the landscape in relation to remotely-sensed forest and landscape structural features. To determine whether snakes selectively forage in areas with higher prey availability, we projected the estimated prey spatial relationships across the landscape and modeled their overlap of occurrence with observed timber rattlesnake foraging locations.
4. We broadly predicted that prey availability was greatest in mature deciduous forests, but *T*. *striatus* and *Sciurus* spp. exhibited greater spatial heterogeneity compared to *Peromyscus* spp. We also combined predicted species encounter rates to encompass a body size gradient in potential prey. The spatial distribution of cumulative small mammal encounters (i.e. overall prey availability), rather than the distribution of any one species, was highly predictive of snake foraging.
5. Timber rattlesnakes appear to select foraging locations where the probability of encountering prey is greatest. Our study provides evidence for fine-scale optimal foraging in a low-energy, ambush predator and offers new insights into drivers of snake foraging and habitat selection.

---

Animal activity patterns are governed by the acquisition of spatially and temporally variable resources from the landscape, such as food, mates, shelter, or hospitable environmental conditions. Successfully procuring food is particularly essential for individual survival, growth, reproduction, and ultimately fitness (Tetzlaff et al., 2017). Therefore, it is important to link the distribution and availability of food with an animal’s space use to better understand the drivers of movement and habitat selection (Heard et al., 2004; Williams et al., 2013).

Given the fitness trade-offs of investing time and/or energy into one behavior instead of another (Beaupre, 2008; Glaudas & Alexander, 2017), optimal foraging theory predicts that predators should forage where they will have the greatest success (i.e. net energy gain; Charnov 1976). More explicitly, the ideal free distribution (IFD) predicts that predators should disperse to patches proportional to their food abundance (Flaxman & Lou, 2009; Williams et al., 2013). Predators can achieve IFD by either tracking prey densities or environmental gradients in prey foraging habitat (Flaxman & Lou, 2009; Kittle et al., 2017). Under game theory predictions, predators are expected to aggregate in areas where prey are abundant at large spatial scales but less precisely match prey distributions at fine scales (Hammond et al., 2012). Accordingly, wolves (Kittle et al., 2017), sea lions (Womble et al., 2009), and snakes (Madsen & Shine, 1996) have all been documented selecting habitats with higher prey availability at broad (regional or macrohabitat) scales.

A complication to understanding drivers of foraging behavior is that habitat selection can be multi-scale and hierarchical (Johnson, 1980; Mayor et al., 2009). Predators can demonstrate hierarchical foraging behavior as a result of multiple scale-dependent processes, such as predation risk or resource availability (McNeill et al., 2020). Conversely, observed foraging patterns can result from predominantly fine-scale resource selection (Harvey & Weatherhead, 2006). The space use of ectotherms is often driven by microhabitat conditions that affect their ability to thermoregulate, avoid predation, and forage (Harvey & Weatherhead, 2006; Sutton et al., 2017). Therefore, snakes may not distribute themselves proportionally to prey availability (i.e. achieve IFD) nor forage optimally if prey-rich patches do not coincide with optimal environmental conditions for thermoregulation (Blouin-Demers & Weatherhead, 2001; Carfagno et al., 2006).

Indeed, some studies have found no evidence of snakes selecting prey-rich areas (Carfagno et al., 2006; Sperry & Weatherhead, 2009; Michael et al., 2014). However, other studies have found mixed or scale-dependent support for prey-mediated habitat selection by snakes (Whitaker & Shine, 2003; Glaudas & Rodríguez-Robles, 2011). Multi-scale studies emphasize the importance of habitat structure coinciding with prey availability for snake habitat selection (Heard et al., 2004; Glaudas & Rodríguez-Robles, 2011). Therefore, whether snakes exhibit IFD characteristics or optimally forage remains unresolved. Investigating the spatial overlap of snakes and their prey is essential to understand potential drivers of foraging behavior and habitat selection.

One hypothesis for the spatial overlap of snakes and their small mammal prey is that similar habitat preferences drive spatial interaction (Blouin-Demers & Weatherhead, 2001). Snakes therefore select habitat based on thermoregulation or other habitat requirements and opportunistically forage, which has been observed in generalist predators such as ratsnakes (*Pantherophis* spp.) and Eastern racers (*Coluber constrictor*; Blouin-Demers & Weatherhead, 2001; Carfagno et al., 2006). Snakes that opportunistically forage may have home ranges containing high prey densities, but they may not exhibit fine-scale selection that maximizes potential prey encounters (Sperry & Weatherhead, 2009). An alternative hypothesis is that the spatial distribution of prey abundance drives snake habitat selection (Blouin-Demers & Weatherhead, 2001). Prey-mediated habitat selection suggests greater alignment of snake space use with prey availability (i.e. demonstrating IFD). This pattern is more likely to be evident in dietary specialists (Madsen & Shine, 1996) or during times of environmental stress such as drought (Whitaker & Shine, 2003).

Although some studies support contrasting hypotheses, not all studies used effective metrics for assessing prey distributions and snake site selection. First, most studies are conducted on a macrohabitat scale, which may not be appropriate when investigating snake habitat selection (Harvey & Weatherhead, 2006). Additionally, researchers typically evaluate prey abundance rather than prey availability. Prey abundance may not equate to prey availability when factors affecting prey detection are not considered (Sperry & Weatherhead, 2009; Reinert et al., 2011). Specifically, prey may be more abundant in some habitat types but more easily detected by the predator in others (i.e. higher catchability; Hopcraft et al., 2005). To our knowledge, no study has estimated prey availability for snakes at a fine scale and assessed prey distributions as a driver of snake foraging site selection.

The paucity of studies examining prey availability at a fine scale may be due to the logistical challenges of determining the exact microhabitats where the predator forages (Glaudas & Rodríguez-Robles, 2011). However, rattlesnake natural history characteristics make them ideal subjects to test hypotheses related to optimal foraging theory. We sought to determine whether foraging site selection of timber rattlesnakes (*Crotalus horridus; hereafter TRS*), is related to the availability of prey on a fine scale.

Timber rattlesnakes are sit-and-wait ambush predators that may wait at a site for many hours to several days (Clark, 2006). They also have a stereotyped foraging posture, in which they orient their head perpendicular to the long axis of a log or other downed wood while maintaining a tight body coil (Reinert et al., 2011). The species’ conspicuous foraging behavior allows for identification of exact foraging sites. In addition, TRS feed almost exclusively on small mammals, primarily shrews (Soricidae), voles (Cricetidae), mice in the genus *Peromyscus,* chipmunks (*Tamias striatus*), and squirrels (primarily *Sciurus carolinensis*; Clark, 2002). This relatively narrow dietary breadth reduces the potential for complex interactive or conflicting relationships between primary prey, alternative prey and TRS foraging preferences (Carfagno et al., 2006).

Our multi-year radio-telemetry study provided a behaviorally and spatially explicit dataset of TRS activity that allowed us to differentiate among behavior-specific site use and account for individual variation in foraging site selection. The primary goals of our study were to define small mammal spatial distributions and their overlap with observed TRS foraging locations at a fine spatial scale to determine whether TRS optimally forage in prey-rich areas. Our approach entailed (1) quantifying small mammal relative availability with widely-distributed camera traps, (2) projecting small mammal encounters across the study area with landscape predictors, and (3) using radio-telemetry-derived TRS behavioral data and the spatially continuous prey encounter surface to assess the predictive strength of prey availability on TRS foraging site selection.

## Methods

### Study Site

We conducted our study within a mixed-use forest landscape (approximately 5,000 ha) in southeastern Ohio. Vinton Furnace Experimental Forest (VFEF) consists primarily of second-growth forests punctuated by early-successional stands managed through various silvicultural and management practices (ODNRF, 2020). Forest communities in the region vary along topographic gradients. Ridgetops and southwestern-facing slopes are dominated by mixed oak (*Quercus* spp.) and hickory (*Carya* spp.) assemblages and shrubby (*Vaccinium* spp.) understory. Northeastern-facing slopes and river bottoms harbor mesophytic taxa such as *Acer rubrum*, *Acer saccharum*, and *Ulmus rubra* (Adams & Matthews, 2019).

### Camera trap design

Timber rattlesnakes hunt along logs (Reinert et al., 1984) and these microhabitats are also used by small mammals as “runways” (Douglass & Reinert, 1982; Fig. 1). To simulate this foraging behavior, we fixed passive infrared game cameras (Moultrie M-888) to metal fence posts approximately one meter above-ground and positioned them directly overlooking the nearest log (> 15 cm diameter) at each site (Fig. 2). We placed a canister with small holes that contained peanut butter under each camera. Our camera deployment protocol allowed us to obtain fine-scale rodent encounter rates, which we considered more informative of prey availability for TRS than representative macrohabitat-scale estimates of rodent abundance (Reinert et al., 2011).

**Figure 1.**
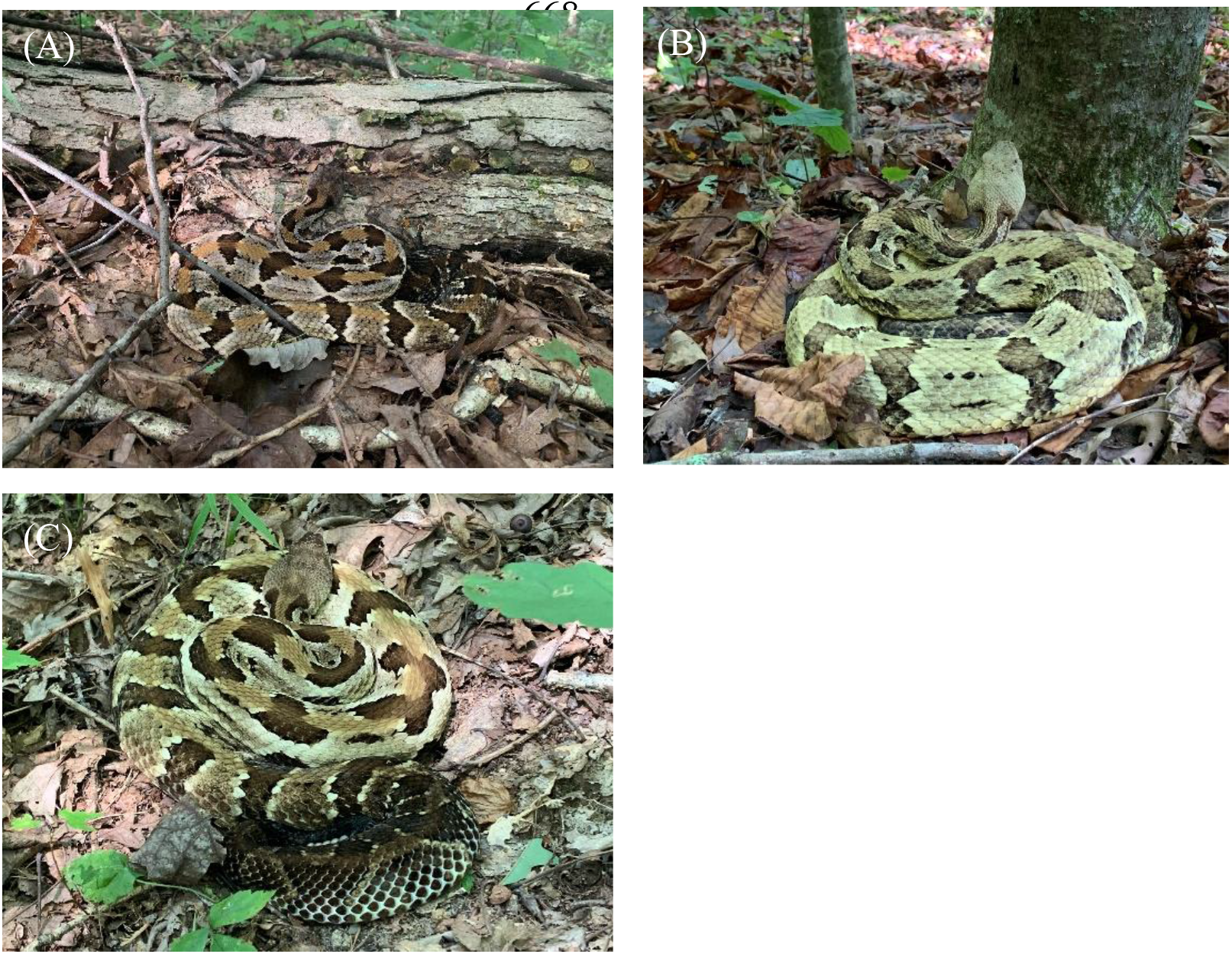
Characteristic ambush posture of timber rattlesnakes (*Crotalus horridus*) used to identify foraging locations, including (A) foraging at logs and downed woody debris, (B) vertical-tree foraging, and (C) non-log foraging along the forest floor. Snakes maintain a tight, “S”-shaped coil regardless of foraging orientation.

**Figure 2.**
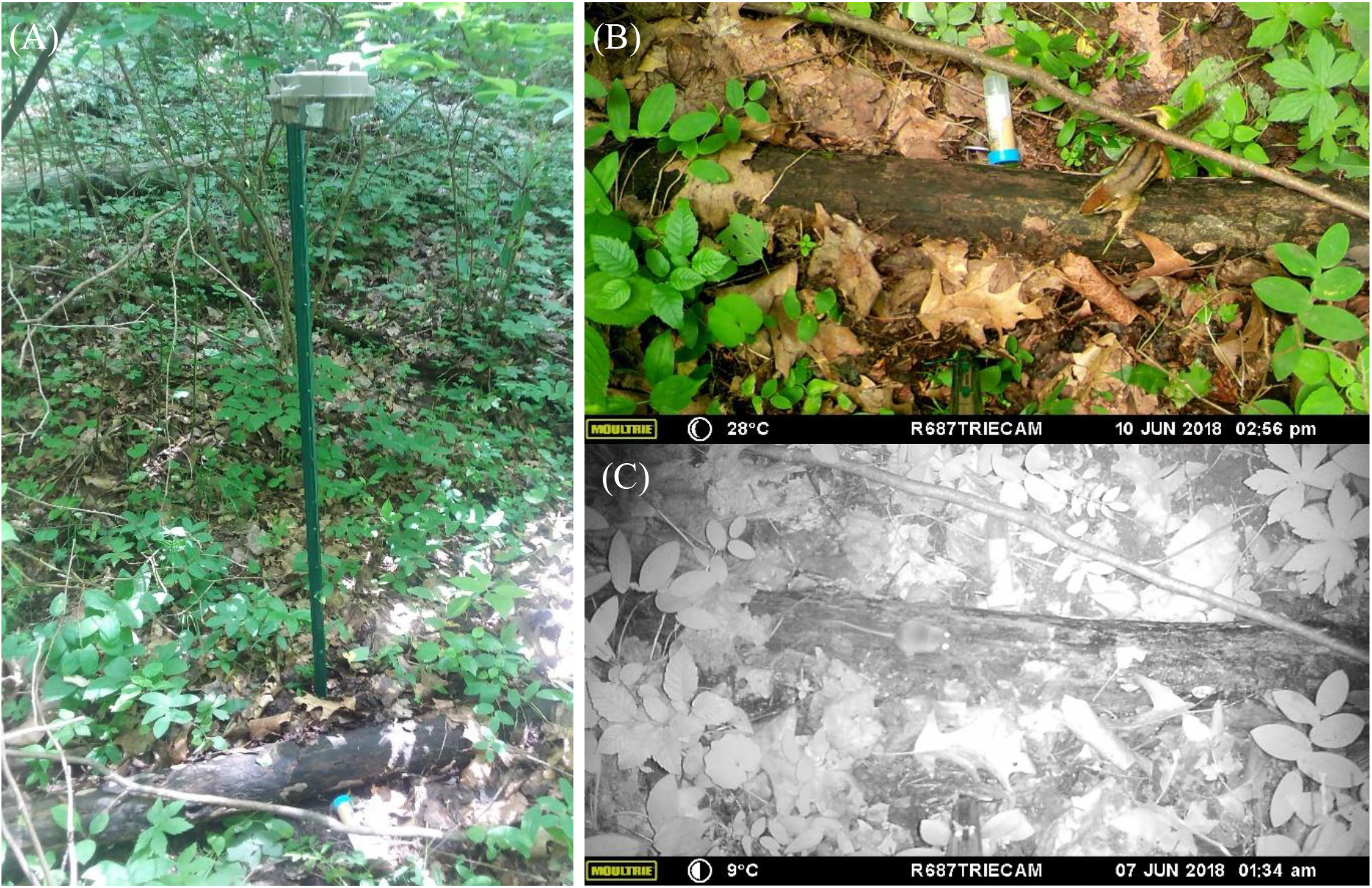
(A) Placement of a camera trap with bait for small mammals one meter above and directly overlooking a log in a mature forest site in southeastern Ohio. (B) *Tamias striatus* and (C) *Peromyscus* spp. captured on camera at this trap site.

We deployed game cameras from 2017–2018 at 242 randomly-chosen, unique sites. We stratified these random points across the dominant macrohabitat types (deciduous forest, pine plantations, clear cuts, and burns) to ensure adequate sampling of each land cover type proportional to its prevalence on the landscape. Accordingly, we sampled more sites from deciduous forests (representing approximately 80% of the landscape) than any other forest type (Table S1). We also set 26 camera traps at previously noted TRS foraging locations. We placed this subset of cameras at observed foraging sites between a day to a few weeks (range 1–86 d; median 15 d) of the snake’s departure from the site.

Game camera active intervals varied by site. We set game cameras at sites for 3–51 days (mean 7.3 d; median 6 d) between 15 June and 13 October 2017 and 4–22 days (mean 8 d; median 6 d) between 24 May and 26 September 2018. We focused our analysis on likely prey items for TRS that were also consistently captured on camera: white-footed/deer mice (*Peromyscus leucopus*/*maniculatus*), eastern chipmunks (*Tamias striatus*), and eastern gray squirrels/fox squirrels (*Sciurus carolinensis/niger*). We monitored occupancy (presence/absence) of each species during observation windows of roughly 12-hr day (approximately 0700–2000 h) and night (approximately 2100–0600 h) periods. We therefore did not track the number of individuals present during each observation period. Because night intervals spanned two dates, we considered small mammals active in the early morning (e.g., before 0600 h) as present in the night interval of the previous date.

### Landscape variables characterizing small mammal distributions

Habitat selection for small mammals, particularly as it relates to forest structural features, is typically assessed with microhabitat and vegetation structural characteristics, such as coarse woody debris and leaf litter cover (Nelson et al., 2019). However, it was not feasible to assess microhabitat features for each camera location and across the landscape. Airborne light detection and ranging (LiDAR) can describe horizontal and vertical vegetation structure across large areas, providing a valuable alternative to the use of intensive field-based methods to assess forest structure (Simonson et al., 2014). Schooler and Zald (2019) demonstrated that LiDAR-derived metrics are effective predictors of small mammal diversity in a temperate mixed-forest community. We therefore used LiDAR and other remotely sensed data to quantify forest structure and predict small mammal occupancy across the landscape.

We described landscape composition and structure at each camera location with 16 land-use, floristic, and topographic variables from fine-scale (5-m) stand-level or remotely sensed data for our study area (Table 1). Stand-level forest management data, including burn history and stand age, reflect active management at VFEF by the Ohio Division of Forestry and U.S. Forest Service over the past 60 years. We derived topographic variables, such as Beers’ aspect (Beers et al., 1966), slope, and elevation from a LiDAR digital terrain model (DTM). To describe forest composition, we considered compositional, multivariate metrics (NMDS1 and NMDS2) that allowed for continuous variation across the landscape. Adams et al., (2019) combined a LiDAR-derived DTM, vegetational plot data, and Landsat 8 OLI imagery to generate floristic gradients for the study area. We sourced the LiDAR-derived DTM from The *Ohio Geographically Referenced Information Program* (*OGRIP; https://ogrip.oit.ohio.gov/Home.aspx*) and Landsat 8 Imagery from the United States Geological Survey (USGS; https://earthexplorer.usgs.gov), and corrected for known timber harvests occurring after data acquisition (Adams & Matthews, 2018; Adams et al., 2019). We tested for multicollinearity among the predictors with Pearson’s correlation coefficient, and no variables were correlated above 0.7. We scaled and centered all continuous variables to have a mean of zero and standard deviation of one.

**Table 1.** Fine-scale (5-m) landscape covariates (n = 16) used to describe small mammal spatial distributions in a mixed-use forest in southeastern Ohio. Summary statistics for each continuous variable provide the mean value (± SD) and range of values across 242 camera trap sites.

| Type | Variable | Description/Interpretation | Summary Statistics<br>(Mean ± SD, range) |
| --- | --- | --- | --- |
| Forest Management | Presence of burns (Burn) | Presence/absence of burned stands (No. camera sites). | Burns = 31<br>Non-burns = 211 |
| | Stand age (Age) | Approximate stand age (years), reflecting forest management activity. | 72 ± 47,<br>4–151 |
| Topography | Beers' transformed aspect index (Beers) | Transforms circular aspect to a range representing xeric southwest aspects to mesic northeast aspects. | 0.95 ± 0.69,<br>0–2 |
| | Elevation (DEM) | LiDAR-derived Digital Terrain Model (m). | 258.4 ± 18.5,<br>203–292 |
| | Slope | Slope (degrees). | 15.3 ± 7.6,<br>0–36.9 |
| | Stream distance (Stream) | Distance to the nearest stream (m). | 154 ± 91,<br>1–381 |
| Multipurpose —<br>Ordination Axes | NMDS axis 1 of floristic variation (NMDS1) | Ordination axis representing a moisture and topographic gradient. Large, negative values represent drier southwestern facing slopes and positive values represent floodplains. | -0.006 ± 0.236,<br>-0.516–0.981 |
| | NMDS axis 2 of floristic variation (NMDS2) | Ordination axis representing a forest structural and successional gradient. Large, positive values are associated with taller canopies and mature forests. | -0.048 ± 0.199,<br>-0.760–0.420 |
| **Vegetation**<br>**Structure &**<br>**Composition** | **Canopy surface height (CHM)** | Canopy height model representing the maximum canopy height (m). | $16.3 \pm 10.7$,<br>0–40 |
| | **MacArthur's Foliage Height Diversity (FHD)** | Shannon's diversity of vegetation hits throughout the vertical profile (3 vertical layers, 0–5, 5–25, and >25 m) of each point location. | $0.70 \pm 0.26$,<br>0–1.04 |
| | **Enhanced vegetation index (EVI)** | Vegetation “greenness” or productivity. | $0.703 \pm 0.109$,<br>0.095–0.878 |
| | **Plant species richness (PSR)** | Number of woody plant taxa. | $14.6 \pm 1.5$,<br>7.8–18.7 |
| | **Overstory density (OVE)** | Amount of overstory (i.e., $\geq 8$-cm DBH) foliage (stems/ha). | $18.5 \pm 24.2$,<br>0–132.2 |
| | **Understory density (UND)** | Amount of understory (<8-cm DBH) foliage (stems/ha). | $164.2 \pm 37.9$,<br>41.9–316.6 |
| | **Tree density (TDE)** | Combined density of overstory and understory size classes of woody plants (stems/ha) | $444.7 \pm 130.5$,<br>56.3–954.6 |
| | **Skewness of LiDAR returns (SKE)** | Metric representing forest succession. Skewness of LiDAR returns is expected to be greater (i.e., longer tail) for mature stands compared to younger stands. | $0.93 \pm 1.18$,<br>-0.06–8.45 |

### Timber rattlesnake radio-telemetry

As part of an ongoing study, we radio-tracked 37 adult TRS (21 males and 16 non-gravid females) between 2016 and 2019 to obtain behavior-specific spatial data (further described in Hoffman et al., 2020, in review). We relocated snakes 1–3 times per week and classified behavioral state (e.g., ecdysis, resting, foraging) upon relocation, resulting in 522 observed foraging locations. We noted foraging locations when snakes exhibited a characteristic “S”-shaped ambush posture: compactly coiled, with head extending past outer coil, and a greater number of anterior directional changes compared to a resting state (Fig. 1; Reinert et al., 1984). We also identified the presumed foraging orientation type—log-oriented, non-log-oriented, or vertical-tree-oriented (Reinert et al., 2011; Goetz et al., 2016). We defined a log-oriented posture as when snakes rested on or faced (within 1-m) a log or fallen branch (Fig. 1A; Reinert et al., 1984). We defined a vertical-tree-oriented posture as when snakes coiled at the base of standing trees, with their heads oriented upwards or facing (within 1-m) a tree (Fig. 1C; Goetz et al., 2016). We considered snakes coiled in ambush on the forest floor but not log or vertical-tree-oriented to be in a non-log-oriented posture (Fig. 1B; Reinert et al., 2011). We found males and non-gravid females in our study equally likely to forage log-oriented (n = 244) as non-log-oriented (n = 239), and to rarely exhibit a vertical-tree-orientation (n = 39). A preliminary analysis showed that foraging orientation type did not affect prey encounter rate for any prey species (Fig. S1). We therefore included all foraging location types in the Snake Foraging Probability models to assess snake foraging spatial concurrence with prey availability.

### Small mammal encounter rate models

We modeled the number of days/nights with a small mammal species’ observation using Bayesian zero-inflated generalized linear models (GLM). We considered a zero-inflated framework because of the coarse sampling of small mammals across our site and resulting overdispersion in counts. Ecological datasets often contain a higher frequency of measured zeros than can be accommodated by standard statistical distributions and can therefore violate the assumptions of these distributions (Martin et al., 2005). Zero-inflated models combine two underlying processes, modeling non-zero counts and true zeros with a Poisson or negative binomial process and the potentially false zeros with a binomial process (*Zi*), generating the probability of measuring a zero in error (Zuur et al., 2009).

We tested the global set of landscape covariates (n = 16), year, an offset of the number of active camera days, and *Zi* covariates (i.e. the binomial “false-zero” process) under negative binomial and Poisson distributions, resulting in 10 candidate global models for each species (Tables S2–S4). We suspected that interannual variation, likely representing acorn mast availability (Clotfelter et al., 2007), or the timing of camera placement during each season (i.e. seasonal fluctuations in small mammal activity patterns) could affect our detection success at a particular location. We therefore accounted for temporal variation in species encounter rates for zero-inflated models with the *Zi* term, using no covariates as a null, median date of camera deployment (modeled as a quadratic function), year, and the additive or interactive combinations of median date and year (Tables S2–S4). We used diagnostic plots to compare each model’s predictions of the mean and variance and selected the global model of best fit. For each small mammal species, a zero-inflated negative binomial model best represented encounters but the selected *Zi* covariates varied by species (Table S5).

We examined model coefficients for their magnitude of effect in each selected global model and removed covariates with no or a negligible effect, removed covariates with >15% posterior distribution overlap with zero, and removed covariates with >20% of the posterior within the Region of Practical Equivalence (ROPE; Piironen & Vehtari, 2017). We considered the resulting species model with the lowest leave-one-out statistic (LOO) and Watanabe-Akaike Information Criterion (WAIC) value to be the most parsimonious.

We projected small mammal spatial relationships across the landscape by using the fitted encounter rate model for each species and the corresponding landscape raster surfaces using the ‘raster’ package in R version 3.6.1 (Hijmans 2020; R Core Team 2020). We generated mean encounter probabilities for each species across the study site at a 10-m resolution for 2017 and 2018. In addition to landscapes of species-specific encounter rates, we considered the dietary breadth of adult TRS, and generated grouped prey landscapes by adding the relevant encounter surfaces together. In one group, we combined mouse and chipmunk encounters (Cumulative MC) because they are most likely to be encountered at logs (Douglass & Reinert, 1982). We also combined mouse, chipmunk and squirrel encounters (Cumulative Prey) to capture the body size gradient in prey selection for adult TRS. We extracted the predicted prey species or prey group encounter rates at every TRS location.

### Snake foraging models

We modeled snake foraging using mixed-effects Bernoulli GLMs with foraging behavior as a binomial function of the spatially-explicit small mammal encounter rates. We included a random effect for individual snakes. We tested models with prey type variations for non-gravid adult females (NGF; n = 16), adult males (n = 21), and the combined adult TRS group (n = 37). We excluded gravid females (n = 10) because they fast during gestation (Reinert et al., 1984).

We did not monitor small mammal spatial distributions for two years (2016 and 2019) that we tracked snakes. Although we recognize the potential for prey fluctuations in density corresponding with acorn mast cycles (Clotfelter et al., 2007), the predictive landscape metrics we considered did not vary over the course of the study. We therefore generalized our findings from 2017–2018 to all observations from our telemetry study. We estimated small mammal encounter rates, comprising mouse, chipmunk, squirrel, and the cumulative prey surfaces (Cumulative MC and Cumulative Prey) for 2017 and 2018, but used the two-year averaged encounter rates for each species or species group to represent prey availability in 2016 and 2019. We report results from 2016–2019 but reference the 2017–2018 subset in model selection tables and when applicable in results (see supplementary material for further details). We tested species-level and cumulative prey models for adults collectively, and non-gravid females and males separately. We considered the foraging models with the lowest LOO and WAIC scores as the best-supported model for each group. We used the ‘brms’ package in R to fit all statistical models (Bürkner, 2017).

## Results

### Prey diversity on camera traps

Across 242 camera sites and a cumulative 1901 trap days and 1662 trap nights, we successfully captured the dominant prey species of TRS. We detected mice at most sites (61% of sites with ≥ 1 detection; Table 2) and the most extensively and frequently (range of 0–17 camera days) of any species. We observed chipmunks and squirrels at fewer sites (< 50% of sites) and less frequently (maximum of 9 and 6 days, respectively; Table 2). In addition to these primary prey items, we also infrequently captured shrews (Soricidae), voles (*Microtus* spp.), and cottontail rabbits (*Sylvilagus floridanus*). We also frequently captured bird species that are potential opportunistic prey sources.

**Table 2.** Camera trap (n = 242) daily detections of potential prey items for timber rattlesnakes (*Crotalus horridus*) in a mixed-use forest in southeastern Ohio from 2017–2018. Bird detections include common woodland residents, such as wood thrushes (*Hylocichla mustelina*), ovenbirds (*Seiurus aurocapilla*), and Carolina wrens (*Thryothorus ludovicianus*).

|  | Sites with<br>≥ 1 Detection | Sites with<br>> 1 Detection | Maximum<br>Detections (Days) |
| --- | --- | --- | --- |
| **Mice**<br>(*Peromyscus* spp.) | 148 (61%) | 86 (36%) | 17 |
| **Eastern chipmunks**<br>(*Tamias striatus*) | 89 (37%) | 48 (20%) | 9 |
| **Squirrels**<br>(*Sciurus* spp.) | 70 (29%) | 15 (6%) | 6 |
| **Shrews** (Soricidae) | 14 (6%) | 2 (0.8%) | 2 |
| **Voles**<br>(*Microtus* spp.) | 1 (0.4%) | 0 | 1 |
| **Eastern cottontails**<br>(*Sylvilagus floridanus*) | 25 (10%) | 8 (3%) | 3 |
| **Birds** | 124 (51%) | 65 (27%) | 11 |

### Small mammal encounter rate models

#### Mice

The most explanatory model for daily mice encounters was a zero-inflated (*Zi* = year) negative binomial model with year, burn history, and stand age (Table S5). Mice were most likely to be encountered in non-burns of a younger age (Table 3). Mice were encountered approximately twice as frequently in 2018 (2.22 mice/day; 95% CI: 1.94–2.55) than 2017 (0.91 mice/day; 95% CI: 0.70–1.16; Fig. 3A). Encounters were lowest in burned stands (Table 3), at 0.53 mice encounters per day compared to approximately 0.91 encounters/day in other stand types (Fig. 3B). Stand age was the weakest predictor of mice encounters (Table 3), but encounters were generally more frequent in younger stands (Fig. 3C). Mice could be encountered across the landscape at a range of 0.32–0.81/day in 2017 and 0.84–1.9/day in 2018.

**Table 3.** Bayesian zero-inflated (*Zi*) negative binomial models of mice (*Peromyscus* spp.), chipmunk (*Tamias striatus*), and squirrel (*Sciurus* spp.) encounter rates across 242 camera sites in a mixed-use forest in southeastern Ohio. We modeled the number of camera days with a species detection, offset by the total number of active camera days, as a function of study year (2017–2018) and remotely-sensed landscape variables (5-m resolution). We report the variables that best described each species’ distribution. Mean coefficient estimates, standard errors (± S.E.) and percentage of the posterior distributions overlapping zero are provided. Refer to Table 1 for further descriptions of covariates.

|  | Mice | Chipmunks | Squirrels |
| --- | --- | --- | --- |
| **Year** | 0.56 (± 0.17)<br><1% | | 1.05 (± 0.31)<br><1% |
| **Burn** | -0.52 (± 0.19)<br><1% | | |
| **Age** | -0.09 (± 0.06)<br>7% | | |
| **Slope** | | 0.19 (± 0.09)<br>3% | |
| **NMDS1** | | | -0.24 (± 0.14)<br>4% |
| **NMDS 2** | | 0.27 (± 0.10)<br><1% | 0.38 (± 0.17)<br>1% |
| **PSR** | | 0.23 (± 0.11)<br>2% | |
| **FHD** | | | -0.41(± 0.22)<br>3% |
| **OVE** | | | 0.3 (± 0.14)<br>2% |
| **UND** | | | -0.4 (± 0.17)<br>1% |
| ***Zi*: (Median camera deployment date)<sup>2</sup>** | | | -1.52 (± 1.16)<br>5% |
| ***Zi*: Year** | -3.56 (± 2.09)<br>1% | -4.02 (± 2.04)<br>0 | 1.21 (± 2.8)<br>30% |

**Figure 3.**
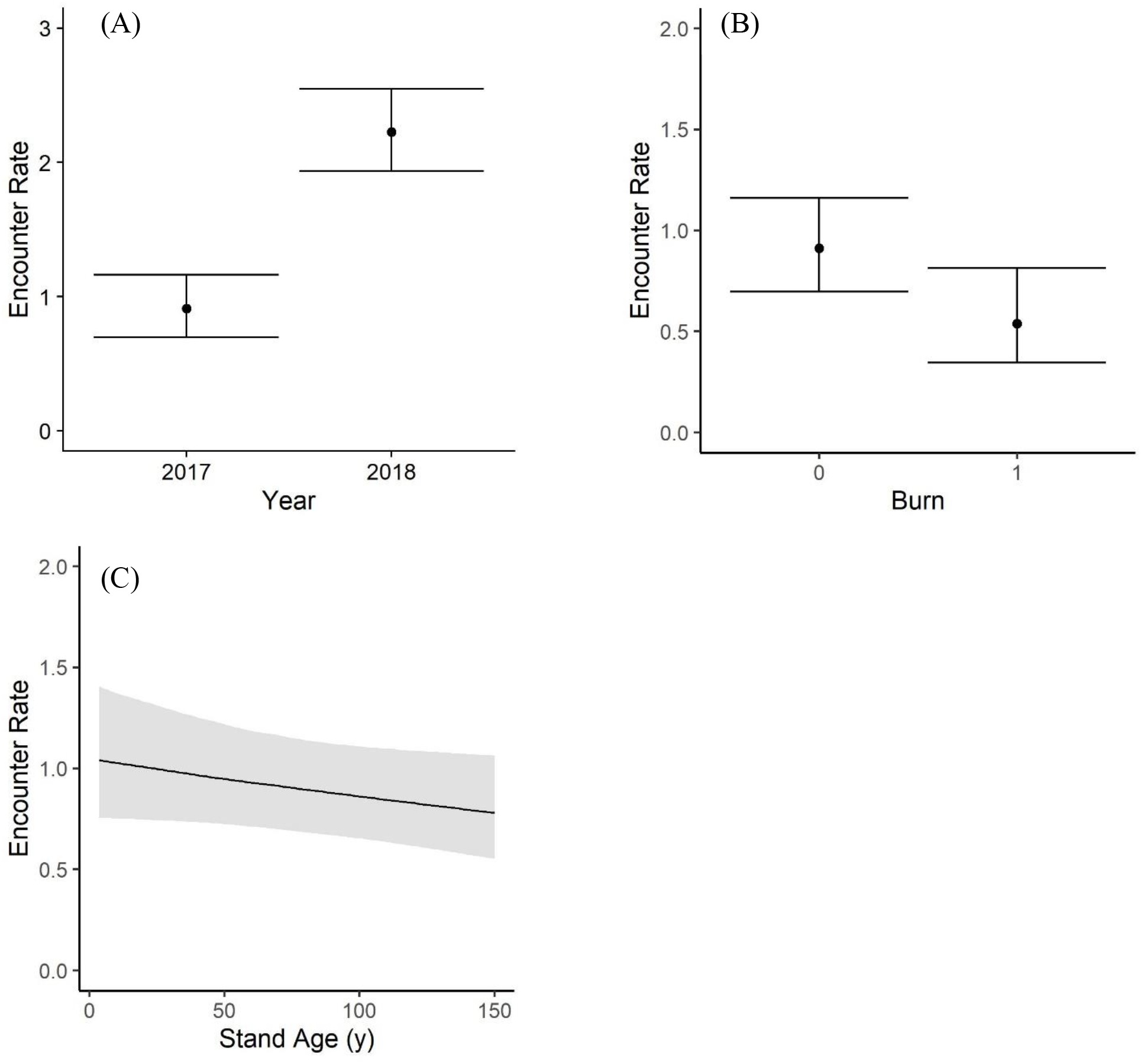
Daily estimated mice (*Peromyscus* spp.) encounter rates from 242 camera trap sites distributed in a mixed-use forest in southeastern Ohio.

#### Chipmunks

The best-fitting model for daily chipmunk (CM) encounters was a zero-inflated (*Zi* = year) negative binomial model with the forest successional gradient (NMDS2), plant species richness (PSR), and slope (Table S5). Chipmunks were most frequently encountered in stands with taller canopies and greater plant richness, and along steeper slopes (Fig. 4). The forest successional gradient was the best landscape predictor (Table 3) of chipmunk encounter rates (Fig. 4A). Encountering a chipmunk would take between 1–2.5 days (mean ~1.5 days) in a more mature forest but between 3–16 days (mean 7 days) in a younger stand. Plant species richness had a similar effect size (mean 0.23 ± 0.11), increasing from 0.14 CM/day (95% CI: 0.04–0.40) at low plant richness to an estimated 0.74 CM/day (95% CI: 0.37–1.48) in a more speciose stand (Fig. 4B). Chipmunks were also encountered more frequently along steeper slopes (Fig. 4C). Chipmunks could be encountered across the landscape at a range of 0.01–1.2/day in 2017 and 0.04–2.8/day in 2018.

**Figure 4.**
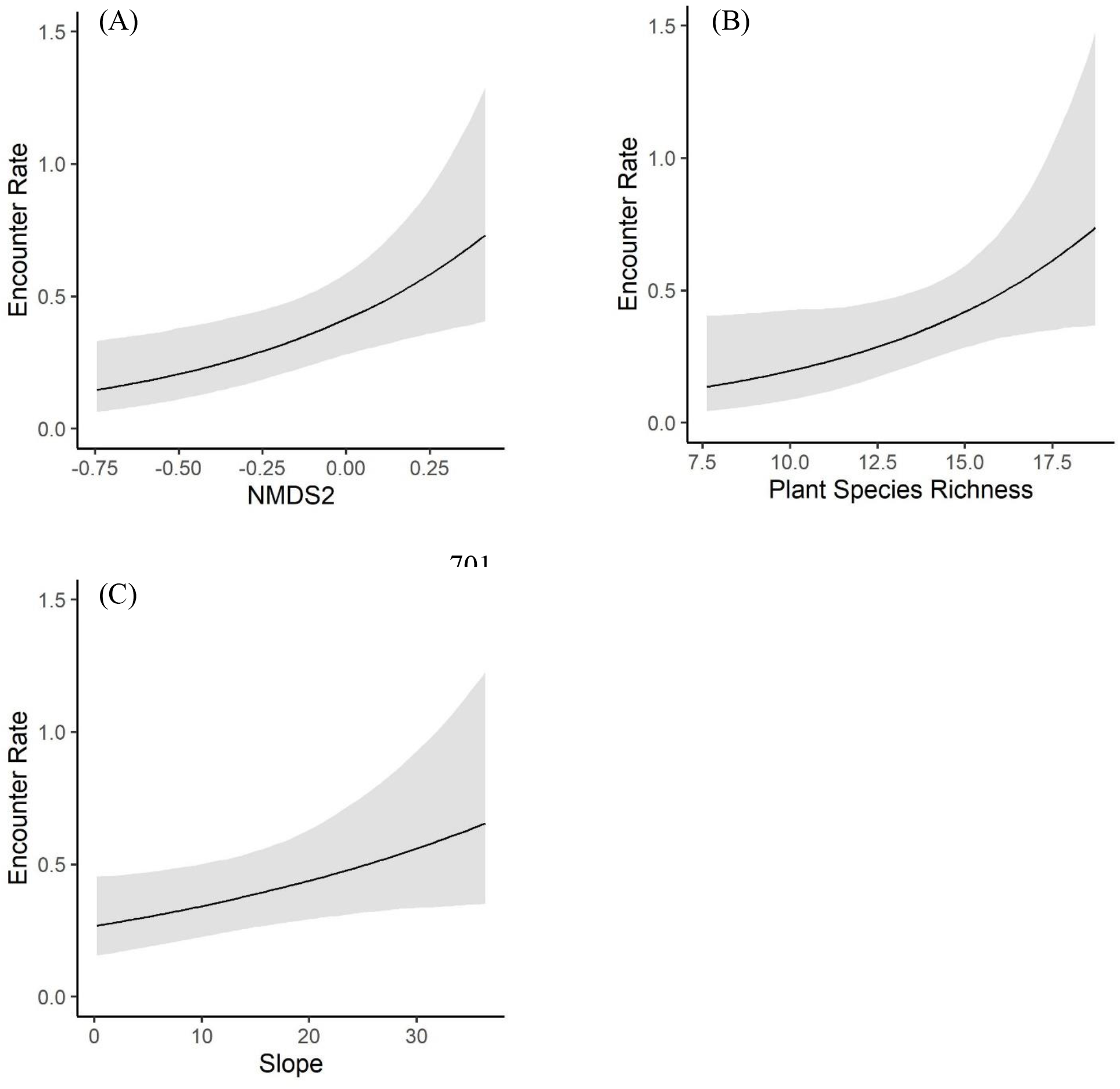
Daily estimated chipmunk (*Tamias striatus*) encounter rates from 242 camera trap sites distributed in a mixed-use forest in southeastern Ohio.

#### Squirrels

The most supported model for squirrel (SQ) encounters was a zero-inflated (*Zi* = median date^2^ + year) negative binomial model with year, foliage height diversity (FHD), the moisture gradient (NMDS1), NMDS2, overstory density, and understory density (Table S5). Squirrels were most frequently encountered in drier areas and stands with taller canopies and greater overstory density, low canopy structural diversity, and low understory density (Table 3). Year had the largest effect on squirrel encounters (1.05 ± SE 0.31), with encounter rate doubling from an estimated 0.16 SQ/day (95% CI: 0.08–0.25) in 2017 to 0.37 SQ/day (95% CI 0.19–0.56) in 2018 (Fig. 5A).

**Figure 5.**
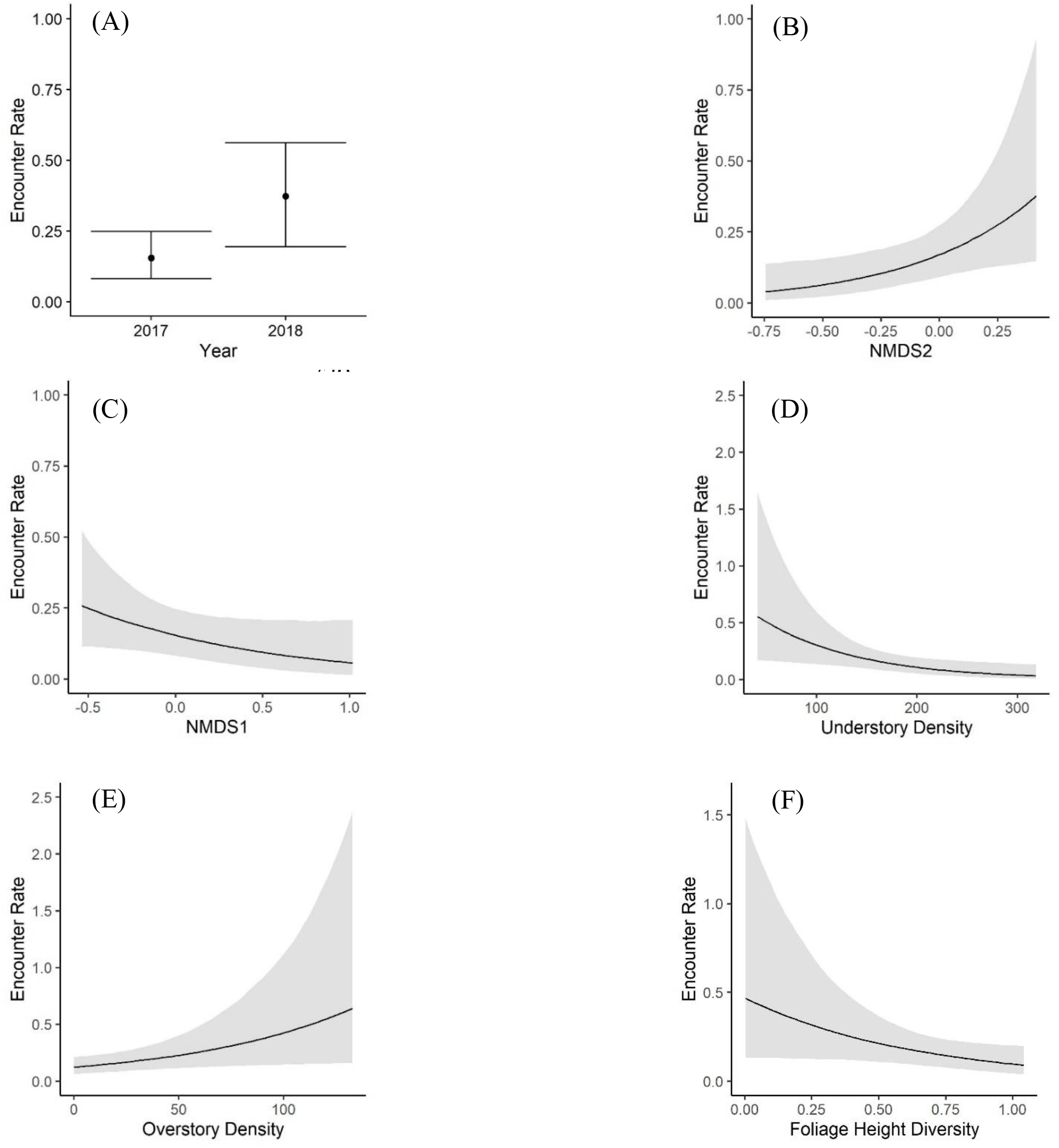
Daily estimated squirrel (*Sciurus* spp.) encounter rates from 242 camera trap sites distributed in a mixed-use forest in southeastern Ohio.

The forest succession gradient (NMDS2) was the best landscape predictor of squirrel encounter rates, with squirrels ten times more frequently encountered in forest stands with taller canopies (0.38 encounters/day, 95% CI: 0.15–0.93) compared to early seral stands (Fig. 5B). Squirrels could be encountered within 1–7 days (mean 3 days) in a mature stand versus rarely, if at all, in the youngest stands with low canopy height. Squirrel encounters were negatively associated with NMDS1, suggesting the species’ preference for drier, southwestern-facing slopes that harbor nut-bearing trees (Fig. 5C). Squirrel encounter rate drastically declined with greater understory density, from 0.55 SQ/day (95% CI: 0.17–1.64) in sparse understories to minimal encounters (0.03 SQ/day; 95% CI: 0.01–0.13) in highly vegetated understories (Fig. 5D). Squirrels were also encountered more frequently at sites with greater overstory density (Fig. 5E). Additionally, squirrels were negatively associated with foliage height diversity (FHD; Fig. 5F), suggesting a preference for forests stands of similar height and age (Aber, 1979). Squirrels could be encountered across the landscape at a range of 0.004–1.29/day in 2017 and 0.01–3.2/day in 2018.

### Foraging Probability Models

The Cumulative Prey landscape, representing the overlay of mice, chipmunk, and squirrel daily encounters, was a strong, well-supported predictor (1.09 ± SE 0.09) of adult TRS foraging probability (Table S6; Table S7). The probability of snake foraging increased sharply with predicted prey encounter rates (Fig. 6). Snake foraging probability increased from a minimum of 0.06 (95% CI: 0.04–0.08) associated with an estimated mean prey encounter rate of 0.59 prey/day to 0.69 (95% CI: 0.61–0.76) at an estimated 3.83 prey/day (Fig. 6).

**Figure 6.**
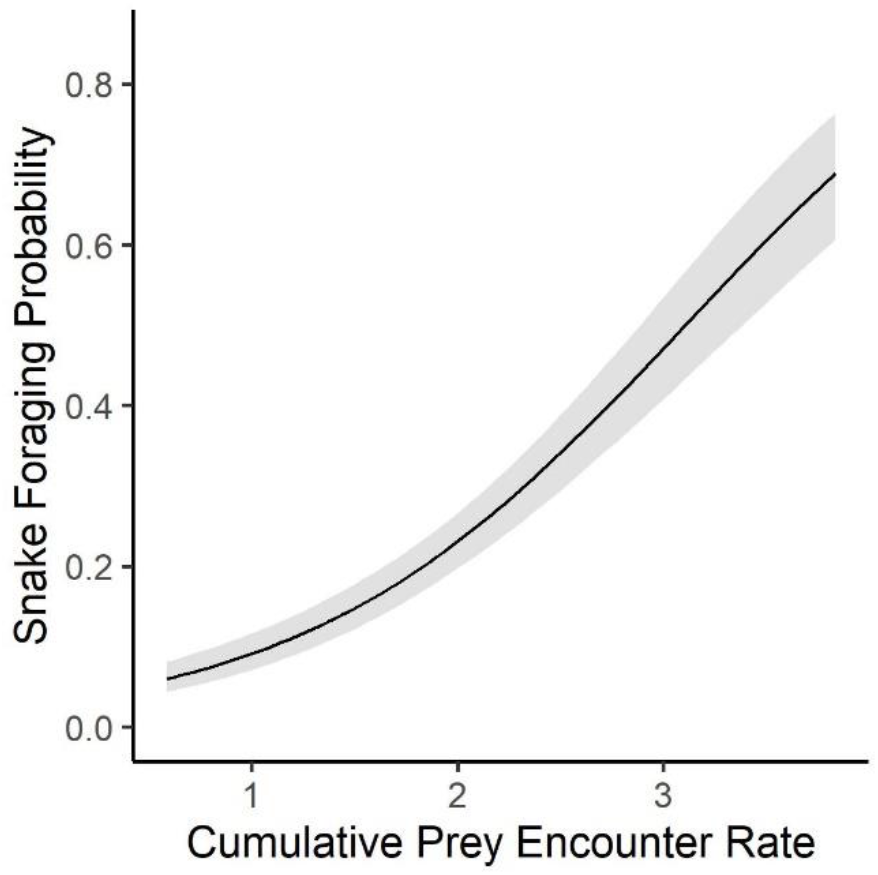
Foraging probability of 37 adult timber rattlesnakes (*Crotalus horridus*) in a mixed-use forest in southeastern Ohio predicted by the cumulative contribution of estimated site-specific, daily encounters with mice (*Peromyscus* spp.), chipmunks (*Tamias striatus*), and squirrels (*Sciurus* spp.) from 2016–2019.

Cumulative Prey was also the best predictor (1.05 ± SE 0.15) for non-gravid females in both year groupings (Table S6). Female foraging probability increased from 0.07 (95% CI: 0.05– 0.11) at low estimated prey encounters (0.63 prey/day) to 0.7 (95% CI: 0.55–0.80) at the highest predicted prey encounter rate (3.8 prey/day; Fig. 7A). In terms of individual prey species, the best-supported species-level model for females included encounter rates for mice (1.39 ± SE 0.22) and squirrels (1.73 ± SE 0.72; Table S7), but with a modest increase in foraging probability associated with mice encounters only (Fig. 7B) and significant uncertainty around the effects of squirrel encounters only (Fig. 7C).

**Figure 7.**
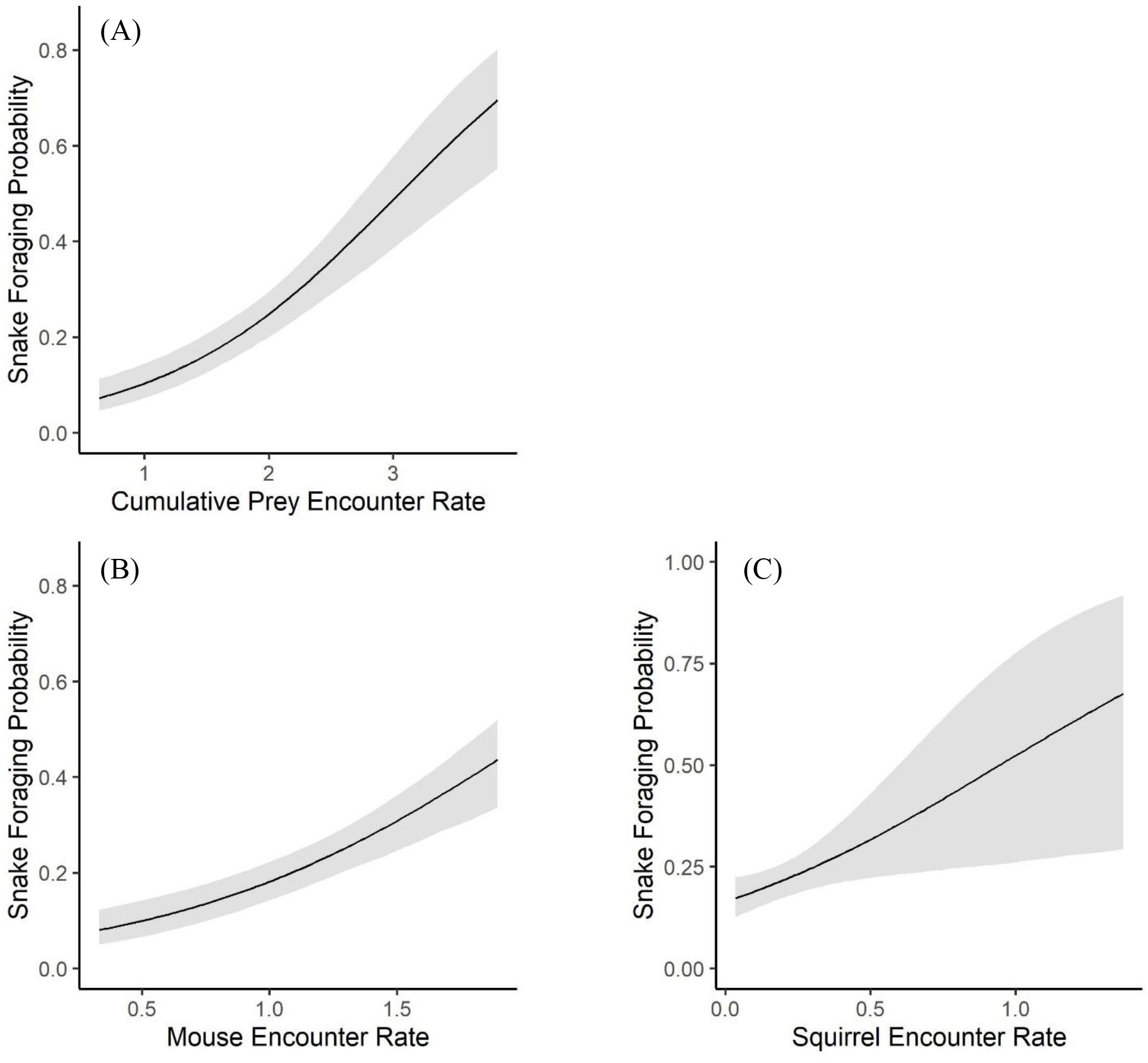
Adult female (n = 16) timber rattlesnake (*Crotalus horridus*) foraging probabilities in a mixed-use forest in southeastern Ohio predicted by prey encounter rates from 2016-2019. A) Cumulative Prey encounter rate, representing the additive combination of mouse (*Peromyscus* spp.), chipmunk (*Tamias striatus*), and squirrel (*Sciurus* spp.) site-specific encounter rates; B) mice-specific encounter rate; and C) squirrel-specific encounter rate.

For males, the species-level foraging model was marginally more supported in 2016– 2019 and the Cumulative Prey model was better-supported in 2017–2018 (Table S6). There was greater uncertainty around species-level effects on male foraging, particularly for squirrels (Table S7). The distribution of mice (0.75 ± SE 0.20) was least predictive of male foraging; foraging probability increased marginally to 0.3 (95% CI: 0.22–0.38) at the greatest predicted encounter rate (1.90 mice/day; Fig. 8A). Predicted chipmunk (1.09 ± SE 0.35) and squirrel (2.9 ± SE 0.77) encounters had the greatest effects on male foraging (Table S7). Male foraging probability increased to a maximum of 0.51 (95% CI: 0.29–0.72) at the highest chipmunk encounter rate of 1.81 CH/day (Fig. 8B). For squirrel encounters, male foraging probability increased sharply to a maximum of 0.62 (95% CI: 0.36–0.83) at high predicted encounters (0.87 SQ/day; Fig. 8C). Estimated Cumulative Prey encounters also had a strong effect (1.14 ± SE 0.11) on male foraging probability, similar to trends observed across all adults (Table S7; Fig. 8D).

**Figure 8.**
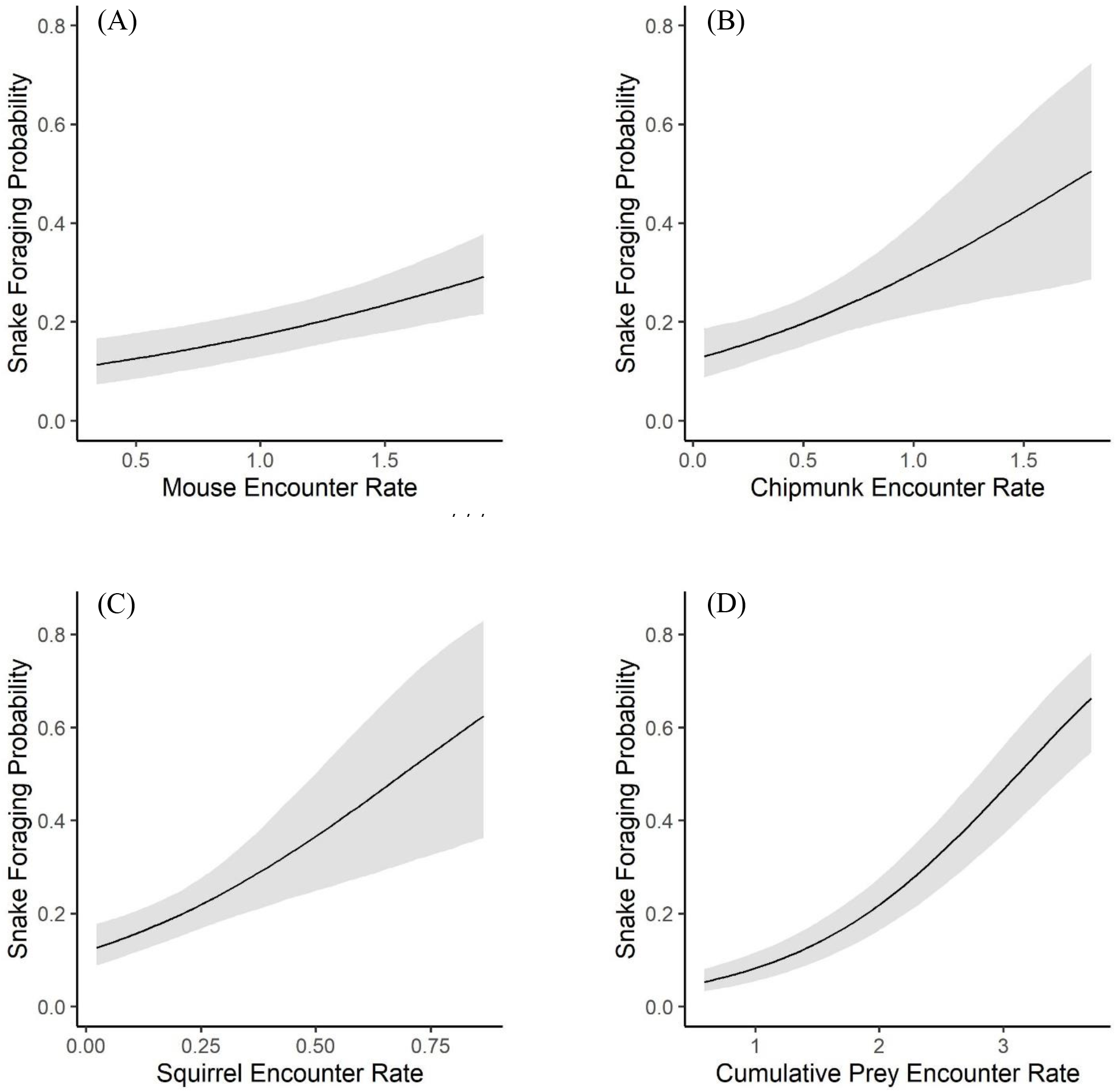
Adult male (n = 21) timber rattlesnake (*Crotalus horridus*) foraging probabilities in a mixed-use forest in southeastern Ohio predicted by prey encounter rates from 2016-2019. Prey-specific foraging models include encounters with A) mice (*Peromyscus* spp.), B) chipmunks (*Tamias striatus*), and C) squirrels (*Sciurus* spp.). D) Cumulative Prey encounter rate, representing the additive combination of mouse, chipmunk, and squirrel site-specific encounter rates.

## Discussion

Because ectotherms have reduced demands for regular, frequent foraging and many snakes in particular can be low-energy specialists (Glaudas & Alexander, 2017), prey distribution and availability may be considered unlikely proximate influences on habitat selection (Heard et al., 2004; Carfagno et al., 2006). However, we found that total prey ‘availability’ (measured as cumulative daily prey encounter rates), rather than any one prey type, was overall the best predictor of TRS foraging. Our results suggest that TRS may forage optimally and preferably forage in prey-rich areas according with IFD. Further, our study supports that TRS may attune to fine-scale differences in prey availability despite specializing on common woodland rodents that are generally thought to be widespread.

Previous studies have found support for some overlap in snake and prey distributions across a single, typically macrohabitat scale. The effects of temporal and/or spatial heterogeneity in prey densities on predator habitat selection is also more straightforward to describe on a broader scale because one can estimate prey abundance/availability and describe snake habitat use within specific habitat types (Glaudas & Rodríguez-Robles, 2011). However, prey may be more abundant in some habitats but more easily detected by the predator in others (i.e. “higher catchability”) due to a lack of cover or camouflage or changes to predator avoidance behavior by prey (Hopcraft et al., 2005). For example, TRS in an agricultural landscape frequently foraged in fields that harbored lower densities of small mammals than surrounding woodlands, likely as a result of increased prey catchability (Wittenberg 2012). Because of the use of both ‘prey availability’ and ‘prey abundance’ interchangeably in the literature, we will hereafter use prey availability to refer to both but will make note of the context in which they are used when possible.

Other studies have shown that snake home ranges generally contain a high proportion of habitat preferred by rodents (i.e. correspondingly high rodent densities), but snakes do not exhibit site selection that would maximize small mammal encounters (Sperry & Weatherhead, 2009; Glaudas & Rodríguez-Robles, 2011; but see Baxley & Qualls, 2009). We expect that we observed a robust, positive association between prey availability and snake foraging on a fine scale partly because we could distinguish foraging site selection from other distinct behaviors shaping site use. Previous snake telemetry studies accounting for prey availability have either aggregated all snake relocations to compare seasonally within home ranges (e.g., Baxley & Qualls, 2009; Sperry & Weatherhead, 2009; Michael et al., 2014) or against random or non-used locations (Glaudas & Rodríguez-Robles, 2011), rather than accounting for behavior-specific variation in habitat use preferences.

Hoffman et al. (2020) found that site selection associated with foraging, ecdysis, digestion, and gestation in this TRS population could be described by many of the landscape variables used in this study (Table 1) at behavior-specific spatial scales (5–105 m). Foraging was negatively associated with temperature and a landscape moisture gradient (indicating drier soils and oak-dominated areas) and these conditions did not describe site use for other behaviors (Hoffman et al., 2020, in review). Importantly, TRS foraging was associated with cooler temperatures than sites associated with other behavioral states, suggesting foraging behavior may be decoupled from snake thermoregulatory needs (Hoffman et al., 2020, in review). Our findings that TRS foraging is associated with greater prey availability but also suboptimal conditions for thermoregulation demonstrate that snakes preferentially seek out prey-rich areas to forage. Habitat structure may therefore incompletely describe foraging behavior, and the prey landscape is an important additional predictor of ambush site selection (Fig. 6).

Our finding that the Cumulative Prey landscape, instead of any one prey species’ distribution, is strongly predictive of TRS foraging can be understood in the context of snake foraging mode and dietary breadth. First, foraging site selection that maximizes encounters across multiple prey species is likely partly due to the sit-and-wait foraging mode of most viperids (Huey & Pianka, 1981; Reinert et al., 1984). Predators using an ambush strategy to hunt are more likely than widely foraging predators to prey on highly mobile species (i.e. species more likely to be encountered), be non-selective in their prey choices, and therefore consume prey species in proportion to their availability (Huey & Pianka, 1981). Viperid species appear to forage in a two-part process, in which snakes first search the surrounding landscape for a suitable ambush site where prey may be more readily available, and then wait to encounter prey or abandon the site when prey encounters are unlikely (Reinert et al., 1984; Clark, 2006). Clark (2006) monitored foraging TRS with videography and demonstrated that snakes selected ambush sites based on potential contact with multiple prey individuals of the same or a different species, with snakes likely using prey chemical trails for identification of these fine-scale small mammal hotspots. Our results also support that TRS may select ambush sites based on the detection of multiple prey species (Clark, 2004).

Specificity in snake diet also affects the importance of the prey landscape for snake habitat selection. The studies that have most conclusively linked snake space use to the abundance of their prey examined focal snake species which primarily consumed a single prey species (Madsen & Shine, 1996; Heard et al., 2004). With increasing diet generalization, snakes are expected to respond to total prey availability rather than the distribution of any one species (Carfagno et al., 2006). This supports our finding that foraging in TRS, a species that primarily consumes small mammals but does not specialize on any species, positively correlates with the overlapping distributions of multiple potential prey.

The orientation of TRS ambush, such as at log, non-log (i.e. forest floor), or vertical-tree, can suggest but not validate the potential prey species targeted through ambush (Reinert et al., 2011; Goetz et al., 2016). Snakes are more likely to encounter mice and some squirrel species (including *Tamias striatus* and *S. carolinensis*) across fallen logs (Douglass & Reinert, 1982), shrews and voles on the forest floor through leaf litter and vegetation (Reinert et al., 2011), and *S. carolinensis* at standing trees (Goetz et al., 2016). An alternative explanation to TRS prioritizing multiple prey chemical cues in ambush site selection is that by combining site-specific prey encounter rates for multiple prey species, we negated any prey-specific preferences by snakes. However, we do not believe this to be likely because we did not detect a difference in predicted encounters of any prey species or combined prey grouping for observed snake foraging sites among ambush orientations (e.g., snakes foraging at logs were not more likely to encounter mice than in non-log ambush). Our finding of equally available prey opportunities among ambush orientations further supports that prey identity is potentially less significant than overall prey availability during foraging in this population (Supplementary Material).

Although Cumulative Prey emerged as the best-supported model for adults generally, we also found some sex-specific differences in individual prey associations (Table S7). Mice most reliably predicted female foraging (Fig. 7B), while chipmunks best predicted male foraging (Fig. 8B). Timber rattlesnakes exhibit an ontogenetic expansion in diet, with larger snakes (i.e. adult males) able to consume larger prey and a broader diversity of small mammals but still target smaller prey indiscriminately (Clark, 2002; Reinert et al., 2011). We must emphasize however that we did not conduct diet analyses to examine the dietary compositions of snakes in our population, and diet has been shown to vary by population and region (Reinert et al., 2011; Wittenberg, 2012; Goetz et al., 2016). We therefore caution against trying to infer dietary patterns from the spatial overlap of snakes with individual prey species or from observed ambush orientations (Clark, 2006; Reinert et al., 2011).

Cameras detected mice (*Peromyscus* spp.) much more frequently than chipmunks (*Tamias striatus*) or squirrels (*Sciurus* spp.), and accordingly, encounter rates for mice scaled higher overall (Table 2; Fig. 3). Despite the prevalence of mice across our study area, their distribution related very little to that of chipmunks (Pearson’s r = −0.19) and squirrels (Pearson’s r = −0.10). Chipmunks and squirrel distributions were most correlated at the landscape scale (Pearson’s r = 0.3). We captured squirrels on camera more intermittently than other rodents, but they exhibited the most complex landscape-scale spatial relationships. Similar to mice and chipmunks, squirrels preferred forest structural characteristics associated with mature forests, including taller canopies and lower understory density, but uniquely with drier, southwestern-facing slopes associated with oaks (Urban & Swihart 2011).

We primarily considered spatial associations of small mammals, but we also observed temporal shifts in availability (Figs 3A and 5A). Rodent encounter rates greatly increased between the two sampled years (2017–2018), and year was the best predictor in the zero-inflated process models for all species (Table S5). We believe this pattern likely corresponds to the boom-bust mast cycles of oaks (*Quercus* spp.), beeches (*Fagus* spp.), and hickories (*Carya* spp.) during 2016 and 2017 and the associated stimulus of increased food availability on rodent population dynamics during the following year (Clotfelter et al., 2007). Given our rodent encounter rate patterns, we suspect, but cannot confirm, that 2016 was a poor mast year and from observational data, 2017 represented a better than average mast crop, particularly for black oaks (*Quercus velutina*) at the site (R. Snell, pers. comm.). We emphasize that we did not expect these yearly fluctuations to affect snake spatial associations with small mammals because our remotely-sensed landscape and forest structural characteristics did not change over the course of the study.

Although our study provides a unique, fine-scale link between prey and predator space use, there are some limitations to the inferences we can make. First, an important assumption to behavioral observations during radio-telemetry is continuity in behavior. We monitored snakes during the day and assumed that individuals remained in a behavioral state if we relocated them at the same site and they exhibited the same behavioral state (e.g. ambush posture) across multiple relocations. We therefore cannot account for temporal gaps in spatial data, during which behavioral shifts or additional ambush site selection/abandonment and any nocturnal foraging patterns may occur (Clark, 2006).

Our field deployment of camera traps was intended to simulate a snake’s perspective and represent a conceptual test of estimating prey availability for this species. Improvements to our camera trap protocol would need to occur in any future applications, such as improving prey species’ detections and sampling more thoroughly and extensively across habitat types and snake ambush sites specifically. Improved image quality could potentially enhance detection of rarely captured, smaller prey, such as shrews and voles (Table 2). Additionally, we used daily species occupancy to account for individual animals moving around or returning to camera sites within a 12-hr interval. Our prey availability metric underestimates true availability. However, we expect this bias to be consistent across all habitats surveyed, which may not be the case in studies using prey abundance as a proxy of availability (Carfagno et al., 2006; Sperry & Weatherhead, 2009).

We also made indirect spatial links between rodents and snakes as our camera locations do not, for the most part, match known TRS foraging locations. We inferred prey availability at snake ambush sites by projecting small mammal spatial relationships across the landscape, which may incompletely capture the spatial heterogeneity of their distributions. However, we found our remotely-sensed landscape-scale covariates (Table 1) to have moderately strong effects on rodent encounter rates (Table 3). In future studies, we recommend direct quantification of small mammal encounters at observed ambush sites and other selected behavior-specific sites to more precisely link foraging behavior to prey spatial distributions.

### Conclusions

Multiple factors could affect the relationship between prey availability and snake spatial ecology, including prey behavior and habitat use, the spatial scales of study, snake diet and foraging mode, and environmental fluctuations. We recognize that thermal requirements are the underlying determinant of overall habitat use variation in snakes, but prey availability plays a potentially important and underappreciated role in local habitat selection. We found a strong association between TRS foraging site selection and rodent encounter rates. Our results suggest that TRS can detect fine-scale differences in prey availability and spatially distribute themselves accordingly. We demonstrate that optimal foraging theory may be applicable to the habitat selection of a low-energy ambush predator.

## Acknowledgements

We thank the Ohio Division of Wildlife for funding this project. B. Borovicka and the U. S. Forest Service provided invaluable logistical support. We also thank T. Lacina, S. Stevens, B. Hiner, E. Scott, and J. Myers for dedicated field work and data collection.

## Authors’ contributions

A. S. H. & W. E. P. designed methodology; A. S. H, J. L. B, & Z. T. T. collected the data; A. M. T. analyzed the data; A. M. T. led the writing of the manuscript. All authors contributed critically to the drafts and gave final approval for publication.

## Data Availability Statement

Data will be made available from the Figshare Digital Repository.

