## Supplemental Tables and Figures for "Prey-Driven Behavioral Habitat Use in a Low-Energy Ambush Predator"

**Supplementary Material**

Table S1. Landcover types present at VFEF, their relative coverage on the landscape, and their representation in our game camera trap dataset for small mammals (242 total camera trap sites).

| Landcover Class | Landscape Coverage | Camera Coverage |
|---|---|---|
| Mature Deciduous | 80% | 183 (75%) |
| Burns (& partial harvest with burn) | 6% | 31 (13%) |
| Clearcuts | 11% | 23 (10%) |
| Pines | 3% | 5 (2%) |
| **Total Sites** | | 242 |

Table S2. Candidate global models describing mice (*Peromyscus* spp.) encounter rates in a mixed-use forest in southeastern Ohio between 2017 and 2018, including variations of zero-inflated (Zi) Poisson and negative binomial models to account for potential temporal variation in small mammal distributions contributing to an excess of zeros. The global set of predictors described forest structure and composition, landscape topography, and year of study (see Table 2 for further descriptions of covariates). Median Date refers to the median date of a camera's active interval and was modeled as a quadratic. We considered diagnostic plots, leave-one-out cross validation (LOO), and Watanabe-Akaike information criterion (WAIC) to determine the most parsimonious model. The best-supported global model (in bold type) was a zero-inflated negative binomial model with year of study explaining the excess of zeros.

| **Distribution** | **Zi Formula** | **LOO** | **WAIC** |
|---|---|---|---|
| Poisson | None | -6.3 | -6.1 |
| | Median Date + Median Date$^2$ | -6.5 | -6.3 |
| | Median Date + Median Date$^2$ + Year | -2.4 | -2.2 |
| | (Median Date + Median Date$^2$) * Year | -3.2 | -3 |
| | Year | -1.5 | -1.4 |
| Negative Binomial | None | -2 | -2.1 |
| | Median Date + Median Date$^2$ | -2.2 | -2.2 |
| | Median Date + Median Date$^2$ + Year | -0.8 | -0.8 |
| | (Median Date + Median Date$^2$) * Year | -1.3 | -1.2 |
| | **Year** | 0 | 0 |

Table S3. Candidate global models describing chipmunk (*Tamias striatus*) encounter rates in a mixed-use forest in southeastern Ohio between 2017 and 2018, including variations of zero-inflated (Zi) Poisson and negative binomial models to account for potential temporal variation in small mammal distributions contributing to an excess of zeros. The global set of predictors described forest structure and composition, landscape topography, and year of study (see Table 2 for further descriptions of covariates). Median Date refers to the median date of a camera's active interval and was modeled as a quadratic. We considered diagnostic plots, leave-one-out cross validation (LOO), and Watanabe-Akaike information criterion (WAIC) to determine the most parsimonious model. The best-supported global model (in bold type) was a zero-inflated negative binomial model with year of study explaining the excess of zeros.

| **Distribution** | **Zi Formula** | **LOO** | **WAIC** |
|---|---|---|---|
| Poisson | None | -9.7 | -9.3 |
| | Median Date + Median Date$^2$ | -10.1 | -9.7 |
| | Median Date + Median Date$^2$ + Year | -5.4 | -4.8 |
| | (Median Date + Median Date$^2$) * Year | -7.3 | -6.9 |
| | Year | -5.3 | -4.7 |
| Negative Binomial | None | -3.4 | -3.4 |
| | Median Date + Median Date$^2$ | -3.5 | -3.6 |
| | Median Date + Median Date$^2$ + Year | 0 | 0 |
| | (Median Date + Median Date$^2$) * Year | -0.8 | -0.7 |
| | **Year** | -0.1 | -0.2 |

Table S4. Candidate global models describing squirrel (*Sciurus* spp.) encounter rates in a mixed-use forest in southeastern Ohio between 2017 and 2018, including variations of zero-inflated (Zi) Poisson and negative binomial models to account for potential temporal variation in small mammal distributions contributing to an excess of zeros. The global set of predictors described forest structure and composition, landscape topography, and year of study (see Table 2 for further descriptions of covariates). Median Date refers to the median date of a camera's active interval and was modeled as a quadratic. We considered diagnostic plots, leave-one-out cross validation (LOO), and Watanabe-Akaike information criterion (WAIC) to determine the most parsimonious model. The best-supported global model (in bold type) was a zero-inflated negative binomial model with median date of camera deployment and year of study explaining the excess of zeros.

| Distribution | Zi Formula | LOO | WAIC |
|---|---|---|---|
| Poisson | None | -1.6 | -1.9 |
| | Median Date + Median Date$^2$ | -0.3 | -0.3 |
| | Median Date + Median Date$^2$ + Year | 0 | 0 |
| | (Median Date + Median Date$^2$) * Year | -0.2 | -0.2 |
| | Year | -2.1 | -2.4 |
| Negative Binomial | None | -1.4 | -1.4 |
| | Median Date + Median Date$^2$ | -0.4 | -0.2 |
| | **Median Date + Median Date$^2$ + Year** | -0.1 | 0 |
| | (Median Date + Median Date$^2$) * Year | -0.5 | -0.5 |
| | Year | -1.5 | -1.6 |

Table S5. Candidate Bayesian zero-inflated (Zi) negative binomial models for mice, chipmunks, and squirrels describing encounter rates from 242 camera traps in a mixed-use forest in southeastern Ohio. The number of camera days with a species' detection (Counts) is offset by the total number of active camera days (Days). Models were reduced from the global (G) set of covariates characterizing forest structure and composition (k = 13), landscape topography (k = 3), and year of study (2017–2018). Refer to Table 2 for descriptions of each covariate. The zero-inflated process was modeled with year and/or median camera deployment date. The most parsimonious model (in bold type) for each species was determined using diagnostic plots, leave-one-out cross validation (LOO), and Watanabe-Akaike information criterion (WAIC).

| **Species & Candidate Models** | **LOO** | **WAIC** |
|---|---|---|
| **Mice (*Peromyscus* spp.)** | | |
| [G] Counts ~ Year + Burn + Age + Beers + DEM + Slope + Stream + NMDS1 + NMDS2 + CHM + FHD + EVI + PSR + OVE + UND + TDE + SKE + offset(log(Days)),<br>Zi ~ Year | -9.6 | -8.8 |
| [M1] Counts ~ Year + Burn + CHM + OVE + Age + offset(log(Days)),<br>Zi ~ Year | -0.9 | -1 |
| [M2] Counts ~ Year + Burn + CHM + Age + offset(log(Days)),<br>Zi ~ Year | -0.1 | 0 |
| **[M3] Counts ~ Year + Burn + Age + offset(log(Days)),<br>Zi ~ Year** | 0 | -0.1 |
| **Chipmunks (*Tamias striatus*)** | | |
| [G] Counts ~ Year + Burn + Age + Beers + DEM + Slope + Stream + NMDS1 + NMDS2 + CHM + FHD + EVI + PSR + OVE + UND + TDE + SKE + offset(log(Days)),<br>Zi ~ Year | -13.6 | -12.9 |
| [C1] Counts ~ Slope + NMDS2 + PSR + OVE + UND + offset(log(Days)),<br>Zi ~ Year | -1.9 | -1.8 |
| **[C2] Counts ~ Slope + NMDS2 + PSR + offset(log(Days)),<br>Zi ~ Year** | 0 | 0 |
| **Squirrels (*Sciurus* spp.)** | | |
| [G] Counts ~ Year + Burn + Age + Beers + DEM + Slope + Stream + NMDS1 + NMDS2 + CHM + FHD + EVI + PSR + OVE + UND + TDE + SKE + offset(log(Days)),<br>Zi ~ Median Date + Median Date$^2$ + Year | -8.2 | -7.5 |
| [S1] Counts ~ Year + Burn + Beers + NMDS1 + NMDS2 + FHD + PSR + OVE + UND + offset(log(Days)),<br>Zi ~ Median Date + Median Date$^2$ + Year | -0.8 | -0.7 |
| [S2] Counts ~ Year + Burn + NMDS1 + NMDS2 + FHD + PSR + OVE + UND + offset(log(Days)),<br>Zi ~ Median Date + Median Date$^2$ + Year | -0.5 | -0.5 |
| **[S3] Counts ~ Year + NMDS1 + NMDS2 + FHD + OVE + UND + offset(log(Days)),<br>Zi ~ Median Date + Median Date$^2$ + Year** | 0 | 0 |

Table S6. Candidate Bayesian mixed-effects Bernoulli models describing timber rattlesnake (*Crotalus horridus*) foraging as a function of predicted prey encounter rates from a landscape-scale small mammal encounter surface (2017–2018) for a mixed-use forest in southeastern Ohio. Species-level models include predicted daily encounter rates for mice (*Peromyscus* spp.), chipmunks (*Tamias striatus*), and squirrels (*Sciurus* spp.). Cumulative models include the additive daily encounter rate predictions for mice and chipmunks specifically (Cumulative MC) or the contribution of all species (Cumulative Prey). Variation in prey encounter rates and snake identity (Snake), modeled as a random effect, described TRS foraging status (Forage) between 2016–2019. In 2016 and 2019, prey encounter values for each snake location represented the average predicted rate (prey species or species grouping) between 2017 and 2018. We used diagnostic plots and leave-one-out cross validation (LOO) to determine the most parsimonious models, identified as the smallest selection criterion value (in bold type), for combined adult TRS, non-gravid female, and male foraging probabilities among species-level and cumulative models for 2017–2018 and 2016–2019.

| **Candidate Models** | **2017–2018** | **2016–2019** |
|---|---|---|
| **Non-Gravid Females** | n = 16 | n = 16 |
| *Species-level* | | |
| [1a] Forage ~ Mice + Chipmunks + Squirrels + (1\| Snake) | -1.9 | -0.8 |
| [1b] Forage ~ Mice + Squirrels + (1\| Snake) | -1.5 | -0.2 |
| *Cumulative* | | |
| [2] Forage ~ Cumulative MC + (1\| Snake) | -1.6 | -1.1 |
| **[3a] Forage ~ Cumulative Prey + (1\| Snake)** | 0 | 0 |
| **Males** | n = 15 | n = 21 |
| *Species-level* | | |
| **[1a] Forage ~ Mice + Chipmunks + Squirrels + (1\| Snake)** | -0.5 | 0 |
| *Cumulative* | | |
| [2] Forage ~ Cumulative MC + (1\| Snake) | -4.0 | -6.7 |
| **[3a] Forage ~ Cumulative Prey + (1\| Snake)** | 0 | -1.3 |
| **Adults** | n = 31 | n = 37 |
| *Cumulative* | | |
| [2] Forage ~ Cumulative MC + (1\| Snake) | -8 | -6.6 |
| **[3a] Forage ~ Cumulative Prey + (1\| Snake)** | -2.3 | 0 |
| **[3b] Forage ~ Cumulative Prey * Sex + (1\|Snake)** | 0 | -0.4 |

Table S7. Bayesian mixed-effects Bernoulli models of adult timber rattlesnake (*Crotalus horridus*) foraging from 2016–2019, explained by species-level daily rodent encounter rates rom landscape-scale prey encounter surfaces of mice (*Peromyscus* spp.), chipmunks (*Tamias striatus*), and squirrels (*Sciurus* spp.) or cumulative prey encounter rates encompassing all prey species. We tested species-level and Cumulative Prey models for non-gravid females (n = 16) and males (n = 21) separately, and Cumulative Prey models for adults collectively. Mean coefficient estimates, standard errors (S.E.), 95% lower (LCI) and upper (UCI) credible intervals, and percentage of the posterior distributions overlapping zero are provided. Refer to Table 6 for further descriptions of candidate models.

| | **Covariate** | **Estimate** | **S.E.** | **95% LCI** | **95% UCI** | **Overlap Zero (%)** |
|---|---|---|---|---|---|---|
| **Females** | Mice | 1.39 | 0.22 | 0.94 | 1.83 | 0 |
| | Squirrels | 1.73 | 0.72 | 0.34 | 3.12 | 0.75 |
| | Cumulative Prey | 1.05 | 0.15 | 0.76 | 1.34 | 0 |
| **Males** | Mice | 0.75 | 0.20 | 0.38 | 1.15 | 0.03 |
| | Chipmunks | 1.09 | 0.35 | 0.4 | 1.78 | 0.13 |
| | Squirrels | 2.9 | 0.77 | 1.41 | 4.42 | 0.03 |
| | Cumulative Prey | 1.14 | 0.11 | 0.92 | 1.37 | 0 |
| **All Adults** | Cumulative Prey | 1.09 | 0.09 | 0.91 | 1.27 | 0 |

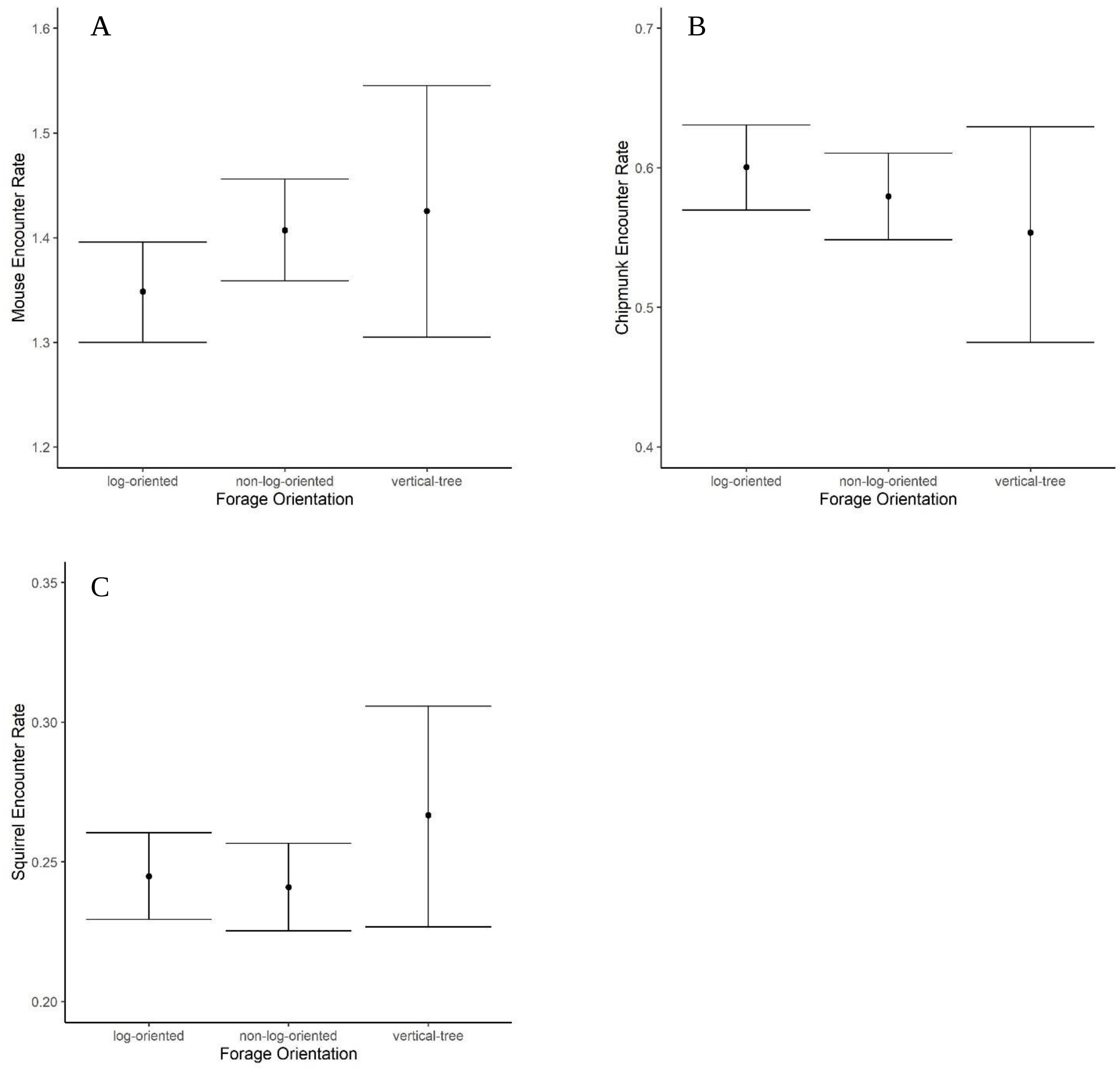


Figure S1. Predicted small mammal prey availability by timber rattlesnake (*Crotalus horridus*) foraging posture orientation. We used a Bayesian multivariate analysis of variance (MANOVA) to examine small mammal prey availability among different foraging orientation types (log-oriented, non-log-oriented, and vertical-tree-oriented) by timber rattlesnakes. We modeled site-specific daily encounter rates of mice (*Peromyscus* spp.), chipmunks (*Tamias striatus*), and squirrels (*Sciurus* spp.) as a function of foraging orientation with the 'brms' package in R (Bürkner, 2017; R Core Team 2020). Log-oriented and non-log oriented were the most commonly observed foraging orientations in our population (n = 244 and 239, respectively) and these foraging orientations exhibited the most similar small mammal associations. There was greater uncertainty around the species-level prey availability of vertical-tree foraging sites due to low observations (n = 39) of this ambush posture in our population. A) Mice encounters were marginally greater (mean: 1.43 mice/day; 95% CI: 1.31–1.55 mice/day) at sites associated with a vertical-tree foraging orientation than non-log-oriented foraging (mean: 1.41 mice/day; 95% CI: 1.36–1.46 mice/day) or log-oriented foraging (mean: 1.35 mice/day; 95% CI: 1.30–1.40 mice/day), but with overlapping credible intervals among all groups. B) Chipmunk (CM) encounters varied little by foraging orientation type. Log-oriented foraging sites yielded the greatest mean chipmunk encounters (0.60 CM/day; 95% CI: 0.57–0.63 CM/day), followed by non-log-oriented (mean: 0.58 CM/day; 95% CI: 0.55–0.61 CM/day) and vertical-tree-oriented foraging (mean: 0.55 CM/day; 95% CI: 0.48–0.63), but with overlapping credible intervals among all groups. C) Predicted squirrel (SQ) encounters were marginally greater (mean = 0.27 SQ/day; 95% CI: 0.23–0.31 SQ/day) at vertical-tree foraging sites, followed by equal encounters at log-oriented (mean: 0.24 SQ/day; 95% CI: 0.23–0.26) and non-log-oriented (mean: 0.24 SQ/day; 95% CI: 0.22–0.26) and overlapping credible intervals among all groups.
